# Ion-mediated condensation controls the mechanics of mitotic chromosomes

**DOI:** 10.1101/2023.04.11.536423

**Authors:** Hannes Witt, Janni Harju, Emma M.J. Chameau, Charlotte M.A. Bruinsma, Tinka V.M. Clement, Christian F. Nielsen, Ian D. Hickson, Erwin J.G. Peterman, Chase P. Broedersz, Gijs J.L. Wuite

## Abstract

During mitosis in eukaryotic cells, mechanical forces generated by the mitotic spindle pull the sister chromatids into the nascent daughter cells. How do mitotic chromosomes achieve the necessary mechanical stiffness and stability to maintain their integrity under these forces? Here, we use optical tweezers to show that ions involved in physiological chromosome condensation are crucial for chromosomal stability, stiffness and viscous dissipation. We combine these experiments with high-salt histone-depletion and theory to show that chromosomal elasticity originates from the chromatin fiber behaving as a flexible polymer, whereas energy dissipation can be explained by interactions between chromatin loops. Taken together, we show how collective properties of mitotic chromosomes, a biomaterial of incredible complexity, emerge from molecular properties, and how they are controlled by the physico-chemical environment.

---

In mitosis, the chromosomes of eukaryotic cells undergo a dramatic metamorphosis, as they are shaped into individualized, rod-like structures with very high density (1). This remarkable transition is realized by the concerted action of several proteins, most particularly condensin I and II, topoisomerase II*α* and the chromokinesin KIF4A (1), leading to the formation of a central protein scaffold, from which the chromatin fiber, composed of DNA and histone complexes, emanates in radial loops (2–4). Simultaneously, chromosomes are compacted by a factor of two (5, 6). Although the name “condensin” suggests that these protein complexes are also responsible for chromosome condensation, rapid depletion of these proteins does not markedly impact chromosome density (7). Instead, it has been shown that the density of mitotic chromosomes is controlled by post-translational modification of histones, in particular through histone deacetylation (8, 9), and is strongly dependent on cations, including Ca^2+^ (10), Mg^2+^ (11–13), and polyamines (13, 14), because of the polyelectrolyte nature of the chromatin fiber (15). Accordingly, polyamine and Mg^2+^ concentrations in the cell are raised during mitosis (12, 16).

Chromosome condensation is followed by attachment of the chromosomes to the mitotic spindle, which segregates the two sister chromatids to the two daughter cells. This is a mechanical process involving forces estimated to be on the order of at least tens of pN (17), indicating that proper biological function of mitotic chromosomes depends on their mechanical properties (18, 19). To characterize these mechanical properties with high precision, we recently introduced an optical tweezers-based approach (20), revealing an intricate nonlinear and visco-elastic response to applied forces. However, the physical origin of these visco-elastic properties has remained elusive. Furthermore, it has recently been shown that chromosome condensation mediated by histone deacetylation is crucial for chromosomal stability (9), leading to the question of how chromosome condensation, chromosomal mechanics, and the biological function of chromosomes are related. A common and powerful approach to understand the mechanics of polymer networks, such as mitotic chromosomes, is to characterize networks prepared at different concentrations or with different polymer modifications, and then use theoretical modelling to determine the underlying physical principles (21–23). This approach is, however, not readily applicable to biomaterials such as mitotic chromosomes, since we have limited control over their composition.

Here, we have overcome this challenge by subjecting optically trapped mitotic chromosomes to two different *in situ* manipulations: ion-mediated chromosome condensation as well as high-salt depletion of histone complexes. We find that stiffness and fluidity of mitotic chromosomes are highly responsive to ionic conditions. By analyzing the change in nonlinear chromosome mechanics upon manipulation, we demonstrate that chromatin in the mitotic chromosome behaves like a flexible polymer. We also characterize chromosomal fluidity and find that it is linked to interactions between chromatin loops.

## *In situ* chromosome manipulation

To enable *in situ* manipulation of mitotic chromosomes, we used an optical tweezers setup integrated with a multi-channel microfluidic flow cell (Fig. 1A). This setup allowed us to attach a chromosome between two optically trapped microbeads via the streptavidin-biotin-interaction (20), and then rapidly and precisely change its physicochemical environment by moving the optical traps with the chromosome to different channels. We then used the optical tweezers to probe the impact of changes in buffer conditions on the structure and mechanics of the trapped chromosome. In this approach, we do not control which one of the different chromosomes is trapped, which introduces substantial intrinsic variability between measurements. Therefore, it is crucial to directly compare the mechanics of the same chromosome before and after manipulation.

**Fig. 1.**
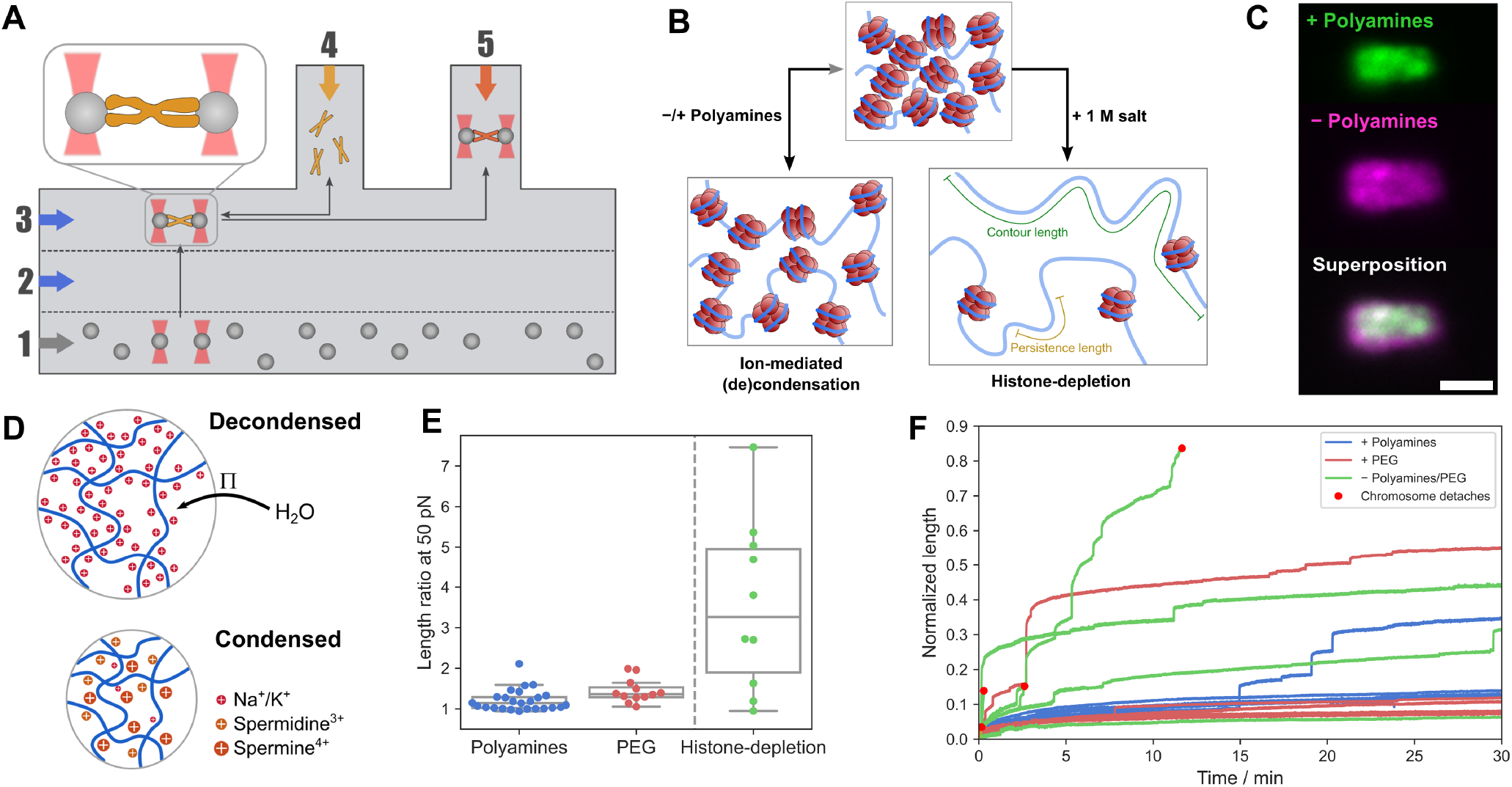
Optical tweezers experiment to examine *in situ* ion-mediated chromosome condensation and histone depletion. (A) Scheme of the experiment, the multichannel flow cell allowed us to maintain different ionic conditions and concentrations in the main channels 1 (also containing streptavidin coated microbeads), 2 and 3 on the one hand and the side channel 5 on the other hand. Chromosomes were introduced into the flow cell via channel 4. (B) Illustration of the two performed manipulations: reversible ion-mediated chromosome condensation and irreversible depletion of histones. The contour length (green) and persistence length (yellow) are indicated in the right panel. (C) Fluorescence images of eGFP-tagged H2B histones of a chromosome in the condensed and decondensed state. Scale bar equals 2 µm. (D) Scheme illustrating the physics of ion-mediated chromosome condensation, the strong negative charge of chromatin needs to be compensated by counterions which lead to an osmotic pressure Π driving water into the chromosome (top). When these counter ions are replaced by polyamines acting as more effective counterions, the chromosome condenses. (E) Box plot of the relative length change caused by polyamines, PEG and histone depletion. To get a robust estimate of the size change, we compared the length of the chromosome *ℓ*(50 pN) at 50 pN before and after changing the solvent conditions. For polyamines, we found a mean length ratio of 121*±*5% (decondensed/condensed, mean*±*standard error of the mean, *N* = 25), for PEG 143*±*8% (*N* = 12), and for histone depletion 355*±*66% (*N* = 10) (F) Normalized chromosome length at a constant force of 250 pN. Without polyamines or PEG, chromosomes frequently elongated stepwise and often disintegrated to the point where they could not support a force of 250 pN (red dots) (for polyamines *N* = 5, for PEG *N* = 4, without polyamines or PEG *N* = 7).

We applied two different manipulations: in addition to changing the ionic conditions to reversibly condense or decondense chromosomes (24), we depleted histone complexes irreversibly by subjecting the chromosomes to high-salt conditions (Fig. 1B) (2). For the first manipulation, we used the physiological, short-chained polyamines spermidine^3+^ and spermine^4+^ (14) (Fig. S1). When a chromosome was moved from polyamine-free buffer to a buffer supplemented with physiological concentrations of spermidine^3+^ (0.5 mM) and spermine^4+^ (0.2 mM), we observed an immediate decrease in chromosome size, which was apparent in force-distance measurements as well as brightfield images and fluorescence images of chromosomes containing eGFP-H2B (Fig. 1C). These changes were mostly reversible when returning to the initial buffer. While polyamine-induced compaction was rapid, decompaction upon depletion of polyamines took several minutes to complete, which might be due to polyamines only slowly detaching from DNA (Fig. S2) (24). Previous work has demonstrated that specific interactions between polyamines and DNA can lead to the formation of DNA-polyamine condensates (25–28). To explain chromosome condensation by polyamines, however, it is not necessary to account for molecular details. Instead, chromosome condensation can be readily explained as shrinking and swelling of a polyelectrolyte gel described by Donnan’s theory (24, 29): Charges in the polyelectrolyte need to be compensated by counterions, leading to a concentration difference of these ions with the bulk of the solvent resulting in an osmotic pressure causing the gel to swell (Fig. 1D, top). Multivalent ions, like polyamines, act as more efficient counterions: by replacing several monovalent cations, they reduce the osmotic pressure inside the chromosome, which in turn leads to shrinking of the polyelectrolyte gel, i.e. chromosome condensation (24, 30) (Fig. 1D, bottom). The density of polyelectrolyte gels can change discontinuously with ion concentration, a phenomenon referred to as a volume phase transition (24, 31).

One prediction of Donnan’s theory is that chromosome swelling is ultimately driven by osmotic pressure and can therefore be counteracted by adding osmolytes, such as long-chained polyethylene glycol (PEG) to the solution (32). Indeed, when we replaced polyamines by PEG-8000 (9% w/v), we still observed efficient chromosome condensation (Fig. 1E), comparable to polyamines, as previously reported (24). Donnan’s theory also predicts that lowering the ionic strength leads to an expansion of the chromosome, while increasing the ionic strength causes further condensation, in agreement with our observations (Fig. S3). Polyamines led to an inversion of the chromosome’s response to the ionic strength since the interaction between polyamines and DNA itself depends on the ionic strength (33, 34).

Notably, we observed that decondensed chromosomes were more fragile than condensed chromosomes (Fig. 1F). When chromosomes were kept at a constant force of 250 pN for 30 minutes in the presence of condensing agents like PEG or polyamines, they showed some viscoelastic creep, but rarely extended by more than 10% (1 out of 5 chromosomes for polyamines, 1 out of 4 chromosomes for PEG). By contrast, in the absence of polyamines or PEG the chromosomes frequently exhibited stepwise elongations. For 4 out of 6 chromosomes analyzed, chromosome integrity was even compromised such that the chromosome was not able to support a force of 250 pN (marked with red dots in Fig. 1F), which was never observed in the presence of polyamines or PEG. This shows that compacting agents like polyamines are crucial for chromosomal integrity.

Thus far, we employed reversible swelling and condensation of the chromosome by modulating the electrostatic and osmotic balance between the chromosome and the surrounding medium. In a second manipulation, we apply high salt concentrations (above 1 M) to irreversibly expand the chromosome. Under these conditions, electrostatic interactions between histone complexes and DNA are weakened to an extent that histones detach from the chromosome, an approach already employed in the seminal work by Paulson and Laemmli (2). Indeed, when we exposed a chromosome to a buffer containing 1 M monovalent salt (80 mM KCl and 920 mM NaCl), we observed an immediate swelling of the chromosome (Fig. S4), which was mostly irreversible when returning to regular buffer (Fig. S5). This length increase was substantially larger than during reversible ion-mediated chromosome condensation (Fig. 1E). When we monitored the presence of histones on the chromosomes by measuring H2B-eGFP fluorescence intensity, we observed that chromosomes were almost completely depleted of histones after roughly 10 minutes (Fig. S6). Surprisingly, chromosomes remained structurally intact following this procedure and were able to withstand forces up to 350 pN, which concurs with reports that mitotic chromosomes can even assemble in the absence of histones (35).

## Origin of elasticity in mitotic chromosomes

The two manipulations introduced here provide complementary insights in the mechanics of mitotic chromosomes. For both manipulations, the chromatin fiber’s persistence length *L*_*p*_ – the length scale beyond which thermal fluctuations significantly bend the polymer (Fig.1B) – is expected to change, since the flexibility of polyelectrolytes is strongly dependent on the charge of the polyelectrolyte and the solvent (36, 37). The contour length *L*_*c*_ of the chromatin fiber in the chromosome (measured along the fiber (Fig. 1B)), is expected to be unchanged during ion-mediated condensation. By contrast, when histones are depleted under high-salt conditions, the contour length of the chromatin fiber increases dramatically as it no longer compacted by histones (2). Thus by compparing the mechanics before and after these manipulations, we can gain complementary insights into the underlying basis of chromosomal mechanics.

First, we considered how the nonlinear chromosome mechanics change during ion-mediated chromosome condensation. To this end, we measured force-displacement curves on the same chromosome in the absence and presence of polyamines or PEG. These curves showed the characteristic nonlinear mechanical response reported previously (20), independent of the condensation state (Fig. 2A). At low forces, the force response was approximately linear with constant stiffness *K*_0_, before stiffening sets in at the critical force *F*_*c*_ and the stiffness increased dramatically. Both linear stiffness *K*_0_ and critical force *F*_*c*_ appeared to increase slightly after chromosome condensation upon polyamine or PEG addition (Fig. 2B, S7, and S8).

**Fig. 2.**
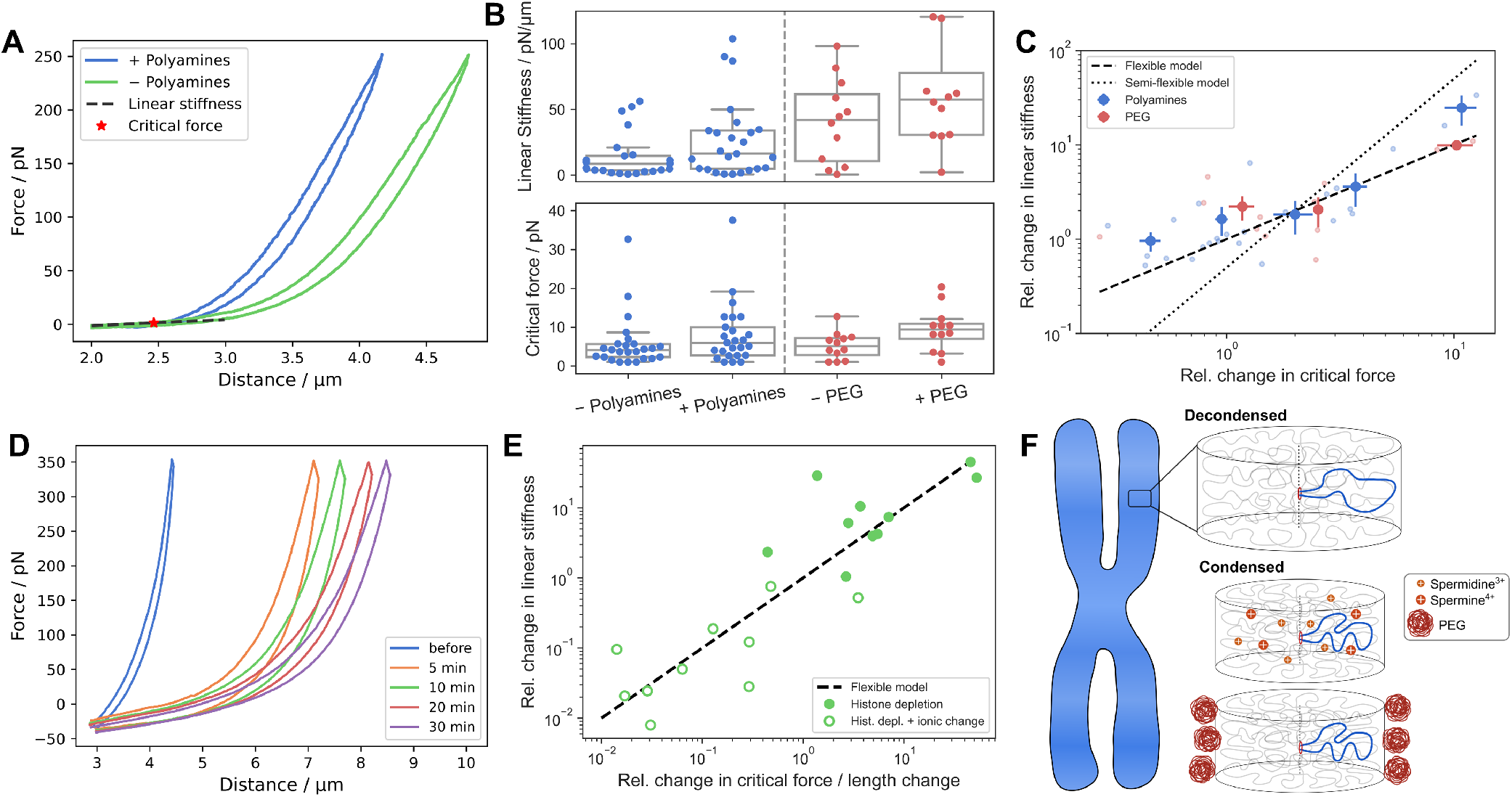
Characterization of nonlinear chromosomal elasticity. (A) Stretch curves of chromosome in the presence and absence of polyamines, indicated are the linear stiffness *K*0 and the critical force *FC* of the decondensed chromosome. (B) Box plots of the linear stiffness and the critical force in the absence or presence of polyamines or PEG (linear stiffness: *p* = 0.19 for polyamines (*N* = 25), *p* = 0.03 for PEG (*N* = 12), critical force: *p* = 0.03 for polyamines, *p* = 0.09 for PEG) (C) Relative change of the linear stiffness 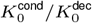 as a function of the change in relative critical force 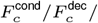 reveals a linear relationship that agrees well with the predictions for a flexible polymer *k*_0_ *∝ F*_*c*_, while it clearly deviates from the prediction for a semiflexible polymer 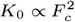. (D) Stretch-curves of a chromosome before histone depletion and after varying times of treatment with 1 M salt, measured in regular buffer containing polyamines. We see a drastic elongation but the general curve shape and the mechanical stability are maintained. (E) Relative change in linear stiffness 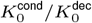 as a function of 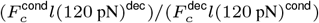 revealing a linear scaling. Filled symbols compare chromosomes before and after histone-depletion, open symbols compare chromosomes before histone-depletion with chromosomes during histone-depletion (while in 1 M salt buffer) (*N* = 10). (F) Scheme illustrating how the mechanical properties of a mitotic chromosome composed of a flexible chromatin fiber depend on the physico-chemical environment.

Importantly, when we analyzed how these parameters changed for each individual chromosome by considering the ratio of the linear stiffness in the condensed and decondensed state 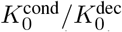 as a function of the ratio of the critical Forces 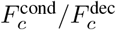, we observed an approximately linear relationship, independent of whether polyamines or PEG were used to condense the chromosome (Fig. 2C). Fitting a power law 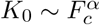 to these data revealed a power-law exponent of *α* = 1.1 ± 0.24. This observation has important implications because this scaling depends on the physical characteristics of the underlying polymer. To that end, we model the chromosome as a single effective worm-like chain (WLC) described by *L*_*p*_ and *L*_*c*_ and consider the relation of linear stiffness *K*_0_ and critical force *F*_*c*_ (see methods). When *L*_*c*_ is constant, as expected for ion-mediated condensation, we expect a linear scaling *K*_0_ ∝ *F*_*c*_ for a flexible polymer (*L*_*p*_ ≪ *L*_*c*_) (38), while for a semi-flexible polymer (*L*_*p*_ ≲ *L*_*c*_), we expect 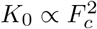 (39, 40). These relations are independent of the persistence length of the polymer, which is dependent on the solvent and the ionic conditions. Comparing these two models to the data, the scaling behavior of chromosomes clearly agrees better with the flexible limit of the WLC model (Fig. 2C).

Next, we consider the changes in nonlinear elasticity upon histone depletion. Again, we found that the general shape of the force-extension curve did not change compared to that of untreated chromosomes, apart from an increase in length and a strong decrease of linear stiffness (Fig. 2D, S5, and S9). This directly showed that, while histones are crucial for chromosome compaction, they do not qualitatively define chromosomal mechanics. During histone depletion, the contour length *L*_*c*_ of chromatin is expected to change dramatically, causing these data to not follow the simple linear scaling between *K*_0_ and *F*_*c*_ (Fig. S10). Thus, the dependency on *L*_*c*_ must be considered (*K*_0_ ∝ *F*_*c*_*/L*_*c*_, see methods). While there is no straightforward way to extract the effective contour length of a chromosome from our data, we assume that the change of the force-dependent chromosome length *ℓ*(*F*) (i.e. the end-to-end distance) upon histone depletion is dominated by the corresponding change in *L*_*c*_, such that 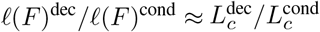 as long as we are sufficiently above the critical force (Fig. 2B). Indeed, when plotting 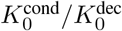 against 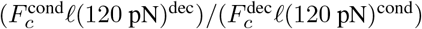, we find a close to linear relation (Fig. 2E, using the length at 120 pN). This not only supports that chromosome mechanics are characterized by flexible polymer behavior but also identifies this polymer as the chromatin fiber.

How do these observations fit our current theoretical understanding of chromosomal mechanics, the Hierarchical Worm-Like Chain (HWLC) model (20)? The HWLC model proposes that mitotic chromosomes are mechanically heterogeneous and behave as a series of nonlinear segments, which are each characterized by a different linear stiffness *k*_0_ and critical force *f*_*c*_. As a result, when a chromosome is stretched, these components stiffen sequentially, giving rise to anomalous stiffening behavior (20), which we still observed independent of chromosome condensation (Fig. S8). This raises the question whether an assembly of heterogeneous elements, i.e. the chromosome, would follow the scaling relations derived above for a single WLC. We found that when all individual components of a HWLC follow a scaling law 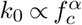, the stiffness of the entire assembly indeed approaches the same scaling 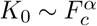 (see methods). Furthermore, for sequential stiffening to occur, the HWLC model requires an exponent *α <* 3*/*2, which excludes semi-flexible WLCs, but is satisfied by chromatin behaving like an assembly of flexible WLCs (*α* = 1), as we argue here.

Taken together, we have demonstrated that chromosomal elasticity originates from the chromatin fiber behaving as a flexible polymer. In this picture, condensation of the chromosome by ions can be understood as a change of the persistence length *L*_*p*_ of chromatin (Fig. 2F), which allows control over the mechanical properties of mitotic chromosomes by modulating their physico-chemical environment, either via more efficient ions like polyamines or osmolytes like PEG.

## Origin of energy dissipation in mitotic chromosomes

Next, we assessed the role of dissipative, viscous contributions in chromosomal mechanics. Therefore, we oscillated the force applied on the chromosome, similar to active microrheology (41). A prestress of 50 pN was applied to the chromosome, before one of the two traps was oscillated with an amplitude of approximately 100 nm at a frequency *ω* of Hz (Fig. 3A) (see methods). This not only allowed us to quantify the elastic stiffness *K*^*′*^(*ω*) from the ratio of the force and distance amplitudes, but also the dissipative, viscous stiffness *K*^*′′*^(*ω*) from the phase shift *ϕ* between force and extension.

**Fig. 3.**
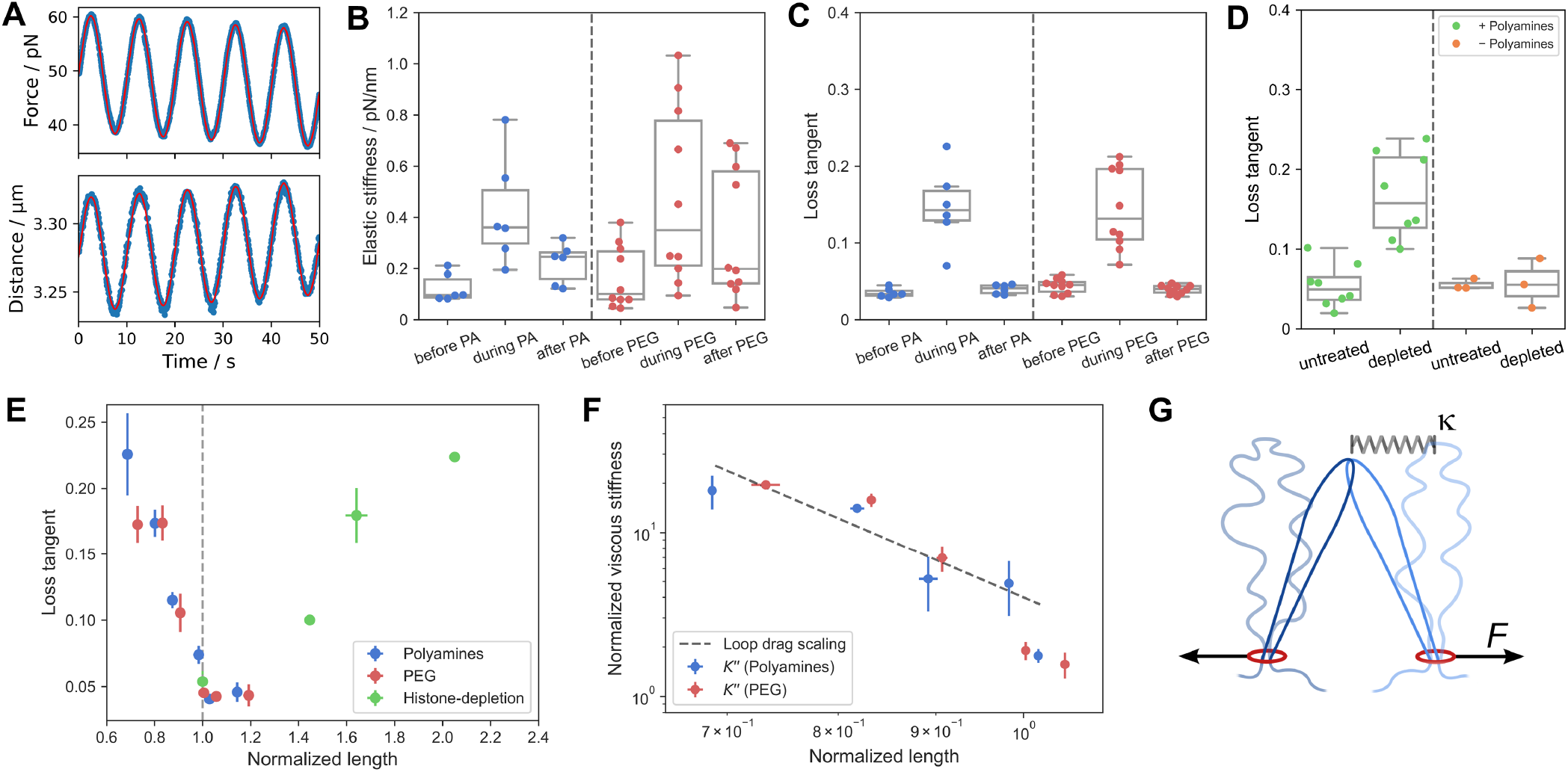
Characterization of chromosomal viscoelasticity. (A) Typical force and distance traces of an oscillatory measurement, data shown in blue, sine fit shown in red. (B) Box plot of the elastic stiffness in the absence and presence of polyamines or PEG. Chromosomes showed strong stiffening from 0.12*±*0.02 pN/nm to 0.42*±*0.08 pN/nm (*p* = 0.02, *N* = 6) for polyamines and from 0.16*±*0.04 pN/nm to 0.48*±*0.1 pN/nm (*p* = 0.003, *N* = 10) for PEG. (C) Box plot of the loss tangent in the absence and presence of polyamines (*p* = 0.003) or PEG (*p* = 0.0004).(D) Box plot of the loss tangent before and after histone depletion in the presence (*p* = 0.0003, *N* = 8) and absence of polyamines (*p* = 0.94, *N* = 3). (E) Loss tangent as a function of length normalized to the length of the decondensed chromosome/the length of the chromosome before histone depletion. (F) The average normalized viscous stiffness as a function of the normalized length, both normalized to the values measured for each respective decondensed chromosome. The dashed line illustrates a scaling of *ℓ*(50 pN)^*−*5^. (G) Proposed mechanism for energy dissipation. A chromosome loop with spring constant *κ* is displaced by a force *F*. Interactions with another loop lead to a drag force extending both loops. The elastic energy stored in the stretched loop is dissipated when they detach.

The elastic stiffness *K*^*′*^ of chromosomes increased strongly when they were condensed with polyamines or PEG (Fig. 3B), as also seen for the linear stiffness (Fig. 2B). Returning the chromosome to polyamine/PEG-free buffer showed that this was mostly reversible on short time scales, in accord with our observations on the reversibility of ion-mediated condensation (Fig. S2). Surprisingly, stiffening of the chromosomes by condensation was accompanied by an increase in fluidity, i.e. the loss tangent tan(*ϕ*) = *K*^*′*^(*ω*)*/K*^*′′*^(*ω*) (expressing the relative importance of elastic and viscous contributions) increased during condensation (Fig. 3C). While in the decondensed state energy dissipation was almost negligible with a loss tangent of only 0.04, during condensation it increased to 0.15 for polyamines and 0.14 for PEG, leading to appreciable energy dissipation under mechanical load. This strong dependency of mechanical properties on a relatively small change in concentration of specific ions means that mitotic chromosomes can be classified as a stimulus-responsive biomaterial (42).

Repeating these experiments on chromosomes before and after histone depletion showed that the fluidity increased strongly after histone depletion in the presence of polyamines, as can be seen from the increase of the loss tangent (Fig. 3D). At first sight, this increase in fluidity as chromosomes expand upon histone-depletion appears to be in contrast to the increase in fluidity as chromosomes condense with polyamines, as both manipulations lead to opposite dependencies of the fluidity on the chromosome size (Fig. 3E). The key mechanical difference between the two manipulations is that in the case of histone depletion the contour length of the chromatin fiber increases. Our results thus indicate that energy dissipation is affected by the length of the chromatin fiber and consequently that the chromatin fiber and its organization are at the origin of the observed differences in fluidity. Furthermore, the observation that the increase in fluidity was identical for polyamines and PEG (Fig. 3E) shows that energy dissipation is not controlled by the charge of the chromatin fiber. It also allows us to rule out mechanisms for energy dissipation based on polyamine-induced attractive interactions between DNA-strands, e.g. by bridging them (26) or by charge inversion (25).

Another clue is the strong density dependency observed upon ion-mediated chromosome condensation: while the length *ℓ*(50 pN) decreased by at most 30% (corresponding to a density increase of a factor ∼3), *K*^*′′*^ increased by a factor of up to 10 (Fig. 3F). This observation suggests a much stronger length dependence than expected for a single polymer. For a Rouse chain, dissipation decreases rather than increases upon condensation (43), whereas the sticky Rouse model, which considers temporary stickers between flexible polymers, predicts a 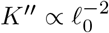 scaling (44), where *ℓ*_0_ = *ℓ*(0 pN) is the rest length of the chromosome, a much weaker dependency than we observed. Stronger scaling of *K*^*′′*^ with concentration has been reported for solutions of flexible polymers (23), but since these models describe isotropic polymer solutions, they seem inappropriate for highly structured chromosomes formed of two long chromatin fibers.

A possible mechanism fitting all qualitative observations is based on the drag the chromatin loops of mitotic chromosomes experience as they interact with each other (Fig. 3G) (45). In this scenario, energy dissipation is expected to be dependent on the average loop size, which is directly related to the contour length, explaining the differences we observed for ion-mediated condensation and histone-depletion. Furthermore, this explanation does not rely on specific interactions between chromatin and condensing molecules, in line with our observation that condensation by polyamines and PEG results in similar mechanical properties.

In the following, we develop a minimal model to demonstrate that this mechanism can also explain the strong dependency of viscosity on the chromosome size during ion-mediated chromosome condensation. We model chromatin loops as springs with stiffness *κ*. When the chromosome deforms, the loops interact, which causes them to extend by a distance ∆*x*, before being released. The resulting dissipated energy scales as *W*_diss_ ∝ *N*_*l*_*κ*∆*x*^2^, where the number of dragged-along loops *N*_*l*_ scales with density and loop sizes. Although chromosomes became softer upon histone-depletion, meaning that *κ* decreases, the increase in loop length could still lead to increased interactions between loops and therefore larger *N*_*l*_ (dependent on the presence of polyamines) and hence increased dissipation.

In the simpler case of chromosome condensation, we can even derive a scaling relation for *W*_diss_. As long as the loop contour length is constant, the number of interacting loops *N*_*l*_ depends only on the density *ρ*, giving us 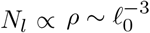 for isotropic chromosome condensation (24). Additionally, condensation results in loops behaving as stiffer springs, which dissipate more energy: for a flexible polymer we have *κ* ∝ ⟨*R*^2^⟩^*−*1^, where ⟨*R*^2^⟩ is the mean squared end-to-end distance. During isotropic condensation, the volume of the chromosome scales with the volume of the loops, such that 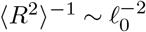. Combining these two relations gives us 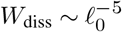 and indeed, although this scaling is derived using the rest length of the chromosome, the imaginary stiffness *K*^*′′*^ at a prestress of 50 pN scales approximately with *ℓ*(50 pN)*−*5 (Fig. 3F).

Taken together, this leads to the following picture of mechanical energy dissipation in mitotic chromosomes: during stretching of the chromosome, the chromatin loops need to rearrange. In the swollen state (in the absence of polyamines or PEG) loops can rearrange readily with little energy dissipation. When the chromosome is condensed, however, the loops collapse, and each loop needs to be dragged through neighboring loops. A similar mechanism has been observed in the friction between polymer brushes, which strongly decreases with the amount of solvent absorbed by the polymers (46, 47) and increases with bristle length (48).

## Conclusion

By combining well-controlled experiments with coarse-grained polymer models, we were able to identify the underlying physical mechanisms for chromosomal elasticity and viscosity even for a system as complex as a mitotic chromosome. We have shown that the material properties of mitotic chromosomes, including density, stability, stiffness and fluidity, are strongly dependent on the physico-chemical properties of their environment, which classifies mitotic chromosomes as a stimuli-responsive biomaterial. This effect, which is inherently due to the polyelectrolyte characteristics of the chromatin fiber and the loop structure of the chromosome, provides cells with a mechanism to control chromosome material properties during the cell cycle by adding or depleting polyamines and other ions. Recent live-cell imaging experiments revealed that stability and proper function of mitotic chromosomes are strongly dependent on histone deacetylation, which like polyamines controls chromosome condensation (9). Our results suggest that the physico-chemical environment of mitotic chromosomes, in particular the presence of polyvalent ions like polyamines, might be equally important for the function, mechanics and stability of mitotic chromosomes.

## ACKNOWLEDGEMENTS

We thank A. Janshoff and A. Biebricher for fruitful discussions.

## Author contributions

H.W., J.H., E.J.G.P., C.P.B., and G.J.L.W. conceptualized the research, H.W., E.M.J.C., and C.M.A.B. performed optical tweezers experiments, H.W. analyzed the data, H.W. and J.H. interpreted the data, T.V.M.C., C.F.N., H.W., and I.D.H. provided isolated chromosomes, J.H., H.W., and C.P.B. developed the model, H.W. and J.H. wrote the initial draft, all authors reviewed and edited the draft.

## Supporting Citations

(49)

