## Supplementary material for "Ion-mediated condensation controls the mechanics of mitotic chromosomes": Methods and Supporting Figures

### Ion-mediated chromosome condensation reveals the origin of viscoelasticity of mitotic chromosomes

**The PDF file includes:**

Materials and Methods
Figs. S1 to S10

### Materials and Methods

#### Buffer compositions

All buffers were made with ultrapure water (MilliQ, Millipore) and filtered (0.2 µm pore size, Whatman) before use.
All buffers contained TRIS (15 mM), EDTA (2 mM), KCl (80 mM), NaCl (20 mM), EGTA (0.5 mM), and Tween20 (0.2%) and were buffered at pH = 8. Polyamine containing buffer additionally contained 0.5 mM Spermidine and 0.2 mM Spermine. Buffer for histone depletion had an increased NaCl concentration (920 mM) to a total concentration of monovalent ions of 1 M. Buffers with doubled salt concentrations contained KCl (160 mM) and NaCl (40 mM), while buffers with a decreased salt concentration contained no KCl or NaCl. PEG-containing buffer was supplemented with 9% w/v PEG8000.

#### Cell culture and chromosome isolation

All cell lines were cultured in High Glucose GlutaMAX DMEM (Gibco) supplemented with 10% fetal bovine serum (FBS, Gibco) and penicillin–streptomycin (Gibco) in a humidified incubator at 37 °C and 5% $CO_2$. For most experiments, chromosomes were isolated from U2OS TRF1-BirA cells with or without H2B-eGFP as described previously *(20)*.
In brief, before isolation, the cell growth medium was supplemented for 48 h with 12.2 mg/L Biotin (Sigma-Aldrich). Cells were stalled in mitosis for 4 h by the addition of 50 ng/mL nocodazole (Sigma-Aldrich). Cells were collected by mitotic shake-off, washed multiple times and then subjected to osmotic swelling in 75 mM KCl and 5 mM Tris-HCl (pH 8.0) for 10 min. Cells were washed and resuspended in polyamine buffer (composition see above), supplemented with Complete mini protease and PhosSTOP phosphatase inhibitor cocktails (Roche). Cells were then lysed in a Dounce homogenizer by 25 strokes with a loosely fitting pestle. After an additional washing step, chromosomes were purified on a glycerol density gradient (60% glycerol (2 mL) and 30% glycerol (2 mL), each in polyamine buffer) at 1750*g* for 30 min. The 60% glycerol fraction contained the purified chromosomes and was stored at -20 °C until use.
For some experiments (see below), chromosomes were isolated from HCT116 TOP2A-mAID H2B-eGFP CDK1as cells infected with lentiviruses introducing TRF1-BirA into the genome (without Auxin addition) as described previously *(20)*.
The experiments for Fig. 1F and 2A-D were performed with chromosomes isolated from U2OS cells, the experiments for Fig. 2D were performed with chromosomes isolated from HCT116 cells, and the experiments for Fig. 1E, 2F and 3 were done with both. We did not observe any differences between their mechanical behavior.

#### Optical Tweezers experiments

The general workflow to study mitotic chromosomes using optical tweezers has been descriped previously *(20)*. Optical tweezers experiments were performed on a commercial optical tweezers setup (C-trap, LUMICKS) or a similarly equipped academic setup described previously *(49)*. In both cases, a 20 W trapping laser (1064 nm, YLR-20-LP, IPG Photonics), a 60x water immersion objective (Plan apo VC NA1.2, Nikon), and multicolor widefield fluorescence imaging were used. A 488 nm excitation laser was used to image both eGFP as well as intercalator dyes. Furthermore, both traps are equipped with a micro-fluidic setup (u-Flux, LUMICKS). The distance between the trapped beads was determined by camera tracking. Forces were determined using a position-sensitive detector and binned to match the frame rate used for distance detection (approx 20 Hz). All experiments were performed using streptavidin-coated polystyrene beads (diameter 4.47 or 4.88 µm, Spherotech). Oscillations of the trap on the academic setup were generated as described previously *(20)*, while oscillations of the commercial setup were enabled by a Bluelake script (LUMICKS).

Trap stiffnesses were calibrated with the thermal noise method before each experiment and were usually around 0.5 pN/nm. Stretch curves were recorded with a pulling velocity of 0.2 µm/s to a maximum force between 250 and 350 pN. Oscillations were performed at 0.1 Hz around a 50 pN prestress. 5 cycles of 6 oscillations each were performed per condition for ion-mediated chromosome condensation, for histone-depletion for each condition one cycle of 10 oscillations was performed. For the stability assay, first a stretch curve was recorded, before the chromosomes were clamped at 250 pN for 30 min or until the chromosome could not maintain the applied force.

Experiments were performed using 5 or 6 channel flow cells. Channel 1 was filled with a given buffer containing in addition the polystyrene beads (5 µL bead stock in 500 µL buffer), while channel 2 and 3 were filled with the same buffer and channel 4 with the same buffer containing chromosomes (20 µL stock in 400 µL buffer). Channel 5 and 6 were then filled with different buffers. Before an experiment was performed under new buffer conditions, the flow cell was flushed for 5 s.

##### Histone depletion

For *in situ* histone depletion, channel 5 of the flow cell was filled with buffer containing a total concentration of monovalent salt of 1 M. Chromosome were first clamped between two beads and then moved to channel 5 for a defined time. Changes in the mechanical properties were assessed by either force-distance curves or by oscillatory measurements. In both cases, one measurement was performed before histone depletion, one measurement during the treatment in channel 5 (towards the end of the specified incubation time), and one measurement after the treatment. For the scaling analysis we not only compared the curves before and after treatment, but also the curves before and during treatment, where we not only see the effect of histone depletion but also of the difference between the regular buffer and the 1 M salt buffer.

#### Data analysis

##### Oscillatory experiments

Oscillatory experiments were analyzed using Matlab and Python scripts. First, the parts of the recorded signal containing oscillations were identified by calculating the cross correlation between the measured distance and the expected oscillation (based on the known number of oscillations and frequency). Therefore, the distance signal was preprocessed by a low pass filter at half the expected oscillation frequency and by detrending the data (although further analysis continued with the raw signal). Next, the parts of the signal containing oscillations was fitted with a sine function. Since force and distance signal are differently measured and processed, it cannot be assumed that the recorded timing is precise. Therefore, in addition to force and distance we also analyzed the bead positions as recorded on the camera, which is in phase with the force acting on the bead and intrinsically synchronized with the distance measure. These three signals (distance, force, and one bead position) were fitted with a sinusoidal function $a_1\cos(2\pi f_1 t) + a_2\sin(2\pi f_2 t) + mt + c$. The sum of cosine and sine yielded more robust fit results and can be transferred to a sine function with a phase shift $A\sin(2\pi ft - \phi)$, with the amplitude $A = \mathrm{sgn}(a_1)\sqrt{(a_1^2 + a_2^2)}$, frequency $f$ and phase $\phi = \arctan(a_2/a_1)$. $\mathrm{sgn}(a_1)$ denotes the sign of the fitted amplitude. The frequency was set at the experimental value and not used as a fit parameter. The complex stiffness was then calculated using the amplitudes of the force and distance oscillations $A_F$ and $A_d$ and the phase of the bead and distance oscillations $\phi_x$ and $\phi_d$ as $K^* = A_F/A_d e^{i(\phi_d - \phi_x)}$.

##### Stretch curves

Length changes during changes in buffer condition were read at 50 pN.
Analysis of force-distance-curves was performed using python scripts using jupyter, the numpy/scipy framework, and matplotlib and seaborn for visualization. In order to calculate the stiffness, force and distance signals were smoothed using a Savitzky-Golay filter from the scipy signal toolbox, with a window length of 81 data points and a 3rd degree polynomial. Then, the stiffness $K$ was calculated by numerical differentiation of force $F$ with respect to distance $d$. To get the critical force a piecewise linear function was fitted to the stiffness as a function of force on a log-log-scale. Therefore, the stiffness was interpolated to a logarithmic force scale, to get evenly spaced data points after taking the logarithm. Then, the logarithms of force $\ln(F)$ and stiffness $\ln(K)$ were calculated and fitted with a piecewise function $y = \ln(K_0)$ for $x \leq \ln(F_c)$ and $y = m \cdot x - m \cdot \ln(F_c) + \ln(K_0)$ for $x > \ln(F_c)$ to determine the initial stiffness $K_0$, the critical force $F_c$, and the stiffening exponent $m$. While this method allows robust determination of the critical force, the initial stiffness provided by this method turned out to be prone to artifacts. Therefore, the linear stiffness was determined by fitting a straight line to the initial part of the force distance curve up to 90% of the critical force. Only data sets where critical force and linear stiffness could be determined unequivocally for the condensed as well as the decondensed chromosome were used for further analysis. This condition was met by 25 out of 42 chromosomes for polyamine-mediated condensation and 13 out of 22 chromosomes for PEG-mediated condensation.
To quantify the scaling behavior of changes in stiffness and critical force, a powerlaw function $ax^b$ was fitted to the data using orthogonal distance regression (scipy.odr) to the individual data points $F_c^{\text{cond}}/F_c^{\text{dec}}$ and $K_0^{\text{cond}}/K_0^{\text{dec}}$. Uncertainties of these data points were calculated by error progression from the uncertainty of the fits used to determine $F_c$ and $K_0$, with a lower bound of 0.25 pN for the absolute error of $F_c$ and 1% for the relative error of $K_0$.
For statistical difference testing, the T-test for related samples was used (function ttest_rel in scipy.stats).

##### Image analysis

Fluorescence images were visualized using imagej. Analysis of the fluorescence intensity of labelled histones was performed in python. Images were loaded using Pylake (LUMICKS), then the background intensity was determined for each image as the most likely intensity of the whole image using a kernel density estimate. Then the total intensity of the background-corrected image was summed up to give the total intensity of fluorescence of labelled histones.

#### Scaling relations

##### Flexible polymer

For the scaling analysis (Fig. 2C, E), we model the chromosome as a single WLC with an effective persistence length $L_p$, and an effective contour length $L_c$, corresponding to the stretched-out length of the chromosome without removal of cross-links. If the effective polymer is flexible ($L_p \ll L_c$), the critical force $F_c$ and the linear stiffness $K_0$ are given by *(38)*

$$F_c = \frac{k_B T}{L_p} \tag{1}$$

and

$$K_0 = \frac{3 k_B T}{2 L_p L_c}, \tag{2}$$

with Boltzmann's constant $k_B$ and temperature $T$. We can combine these two equations to eliminate the dependency on the persistence length $L_p$ to get

$$K_0 = \frac{3F_c}{2L_c}. \tag{3}$$

When we consider relative changes of these parameters between the condensed and the decondensed state (noted by superscript cond and dec, respectively) we find

$$\frac{K_0^{\text{cond}}}{K_0^{\text{dec}}} = \frac{F_c^{\text{cond}} L_c^{\text{dec}}}{F_c^{\text{dec}} L_c^{\text{cond}}}. \tag{4}$$

Since the stretched-out length of a chromosome will be determined by the amount of chromatin between cross-links within the chromosome, $L_c$ will scale with the contour length of the chromatin fiber. Hence, since the contour length of chromatin is constant during condensation, this gives us a prediction for relative changes of critical force $F_c$ and linear stiffness $K_0$ during chromosome condensation:

$$\frac{K_0^{\text{cond}}}{K_0^{\text{dec}}} = \frac{F_c^{\text{cond}}}{F_c^{\text{dec}}}. \tag{5}$$

##### Semiflexible polymer

If the effective contour length of the chromosome is of the order of its persistence length, the critical force $F_c$ and the linear stiffness $K_0$ are given by *(39, 40)*

$$F_c = \frac{k_B T L_p}{L_c^2} \sim \frac{L_p}{L_c^2} \tag{6}$$

and

$$K_0 = \frac{90 k_B T L_p^2}{L_c^4} \sim \frac{L_p^2}{L_c^4}. \tag{7}$$

We can combine the two equations to eliminate the dependency on the persistence length $L_p$ and the effective contour length $L_c$ to find

$$K_0 \sim F_c^2, \tag{8}$$

which directly gives us a predicted scaling between changes of critical force and linear stiffness during chromosome condensation or histone removal:

$$\frac{K_0^{\text{cond}}}{K_0^{\text{dec}}} = \left(\frac{F_c^{\text{cond}}}{F_c^{\text{dec}}}\right)^2. \tag{9}$$

##### HWLC

We now consider a serial assembly of $N$ polymer elements, with critical forces sampled from a power-law distribution $P(f_c) \propto f_c^{-\beta}$, and the linear stiffness scaling as $k_0 \propto f_c^{\alpha}$ *(20)*. Here, $F_c$ and $K_0$ are the critical force and linear stiffness of the whole assembly, i.e. the chromosome, while $f_c$ and $k_0$ are the critical force and linear stiffness of an individual element. The critical force of the assembly will be given by the smallest critical force of its elements; $F_c = f_{c,1}$ for $f_{c,1} < f_{c,2} < \ldots < f_{c,N}$. The assembly's linear response spring coefficient can be expressed as

$$K_0^{-1} = \sum_{i=1}^{N} k_{0,i}^{-1}. \tag{10}$$

By assuming a scaling relation $k_0 \propto f_c^\alpha$ for all elements, we find

$$\frac{F_c^\alpha}{K_0} = \sum_{i=1}^{N} \left(\frac{f_{c,1}}{f_{c,i}}\right)^\alpha. \tag{11}$$

We now claim that the right-hand side of the expression approaches a constant for sufficiently high $N$. To show this, we define a random variable $X(f_c)$ which follows a uniform distribution. The expected separation between two consequent samples, $X(f_{c,i}) - X(f_{c,i+1})$, can then be approximated by a constant proportional to $1/N$. To illustrate, if we sample 10 points uniformly from the interval $(0,10)$ and sort them by magnitude, the distance between two neighbouring samples will be approximately one.

We first take $\beta \to 0$, so that $f_c$ itself is uniformly distributed. We hence have $f_{c,i} \approx f_{c,1} + (i-1)\Delta$, where $\Delta \propto 1/N$. We further note that $f_{c,1} \approx \Delta$, and hence

$$\frac{f_{c,1}}{f_{c,i}} \approx \frac{1}{i}, \tag{12}$$

and Equation 11 hence approaches the Riemann zeta function.

We now take $\beta = 1$. In this case, the variable $\log(f_c)$ is uniformly distributed, and we hence expect that $\log(f_{c,i-1}) - \log(f_{c,i}) \propto 1/N$, and $\frac{f_{c,i-1}}{f_{c,i}} \approx r$ is approximately constant. We then write ratios in Equation 11 as

$$\frac{f_{c,1}}{f_{c,i}} = \frac{f_{c,1}}{f_{c,2}} \frac{f_{c,2}}{f_{c,3}} \dots \frac{f_{c,i-1}}{f_{c,i}} \approx r^{-(i-1)}. \tag{13}$$

Equation 11 then resembles a geometric series with common ratio $r^\alpha$.

Finally, for $\beta \neq 1$, $X(f_c) = f_c^{1-\beta}$ follows a uniform distribution. Rewriting Equation 11 as

$$\frac{F_c^\alpha}{K_0} = \sum_{i=1}^{N} \left(\frac{X(f_{c,1})}{X(f_{c,i})}\right)^{\alpha/(1-\beta)}, \tag{14}$$

we recover the case for a uniformly distributed $f_c$, with $\beta = 0$; the exponent in the sum is simply replaced by $\alpha/(1-\beta)$. We note that the Riemann zeta function converges for exponents $\alpha/(1-\beta) > 1$ and $1 > \beta$ only. Our previous work suggests $1-\beta \approx \alpha - 0.85$, and hence the first condition is satisfied when $\alpha > 0$. Having shown that the right-hand side of Equation 11 is approximately constant when $P(f_c) \propto f_c^{-\beta}$, $\alpha > 1-\beta$ and $\beta \leq 1$, we hence find that for sufficiently high $N$, the stiffness of the HWLC assembly scales as $K_0 \propto F_c^\alpha$ when the individual components follow the same scaling $k_0 \propto f_c^\alpha$.

**Fig. S1**

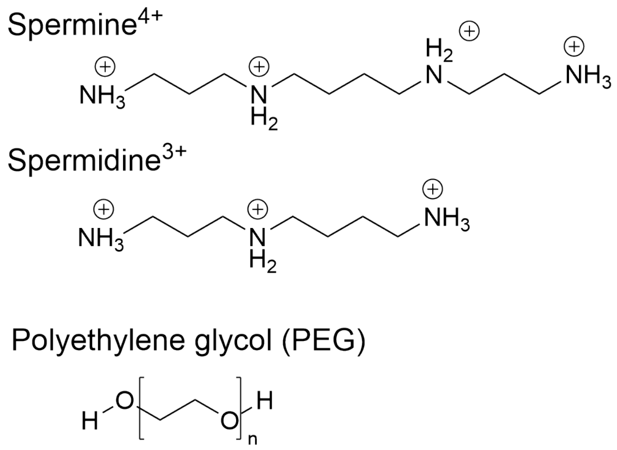


**Figure S1:** Chemical structures of the polyamines spermine$^{4+}$ and spermidine$^{3+}$ and of polyethylene glycol.

### Fig. S2

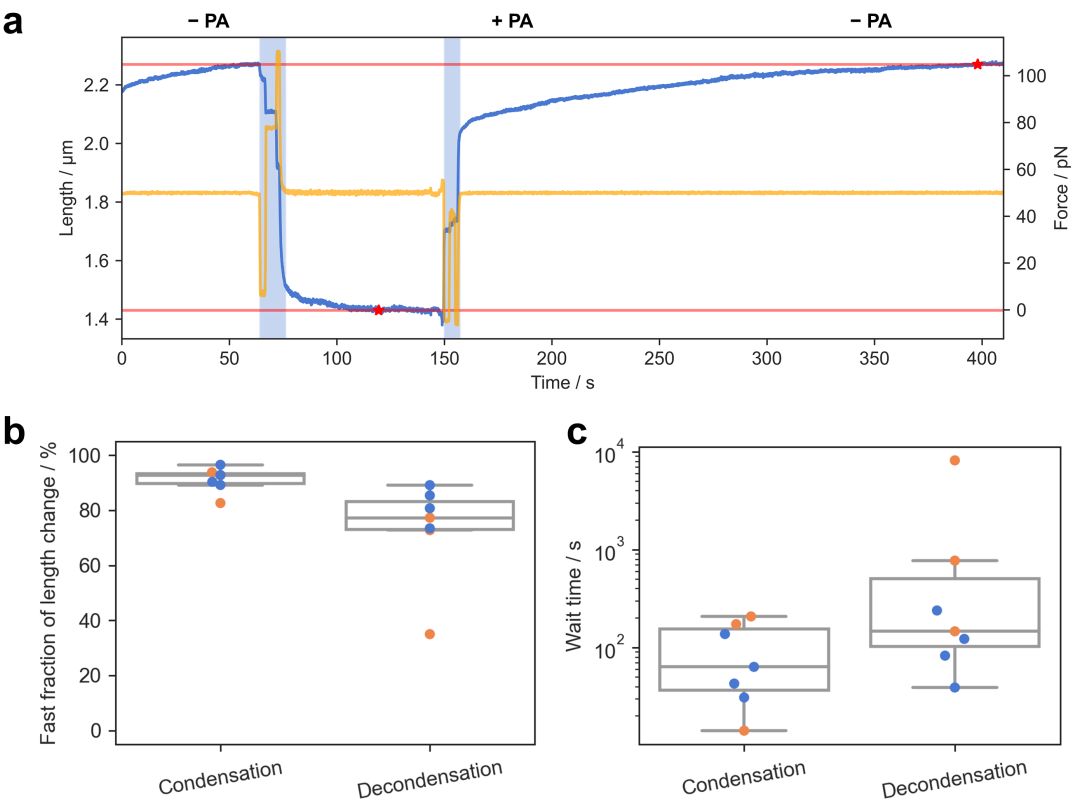


**Figure S2:** Kinetics of chromosome condensation. (a) Exemplary experimental data, a chromosome was clamped at a constant force of 50 pN (orange) and its length was measured (blue). Then the chromosome was moved from polyamine-free buffer to buffer containing polyamines and back (time while moving is shaded blue). We observed an immediate length change followed by a slower creep until a minimal length or the initial length were reached (red star). (b) Percentage of the total length change that was instantaneous given the experimental time resolution (blue dots: 50 pN, orange dots: 20 pN). (c) Wait time until the minimal length or the initial length were reached. Decondensation was much slower than condensation and appears to be force-dependend (blue dots: 50 pN, orange dots: 20 pN).

### Fig. S3

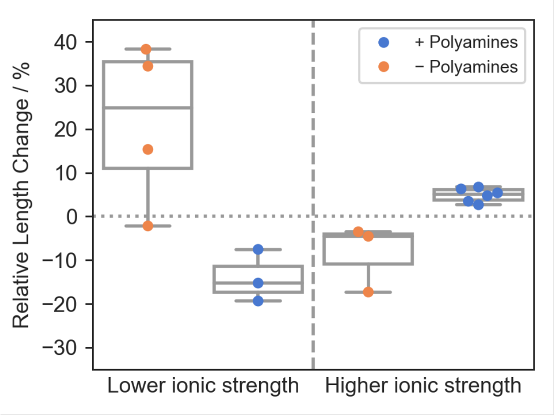


**Figure S3:** Box plot of the length change when a chromosome was moved from a buffer containing 100 mM monovalent salt to a buffer containing 0 mM monovalent salt (left) or 200 mM monovalent salt (right), in the presence and absence of polyamines, respectively. Interestingly, the presence or absence of polyamines qualitatively changed how the chromosomes reacted to changes in ionic strength. Without polyamines, reducing the total monovalent salt concentration from 100 mM to 0 mM leads to an expansion of the chromosome as predicted by Donnan theory, since it increases the osmotic imbalance between the inside and the outside of the chromosome. By contrast, doubling the monovalent salt concentration from 100 mM to 200 mM (still below salt concentrations where irreversible changes occur) leads to a decrease in chromosome size, since the osmotic imbalance is reduced *(24)*. However, when the same changes are performed in the presence of polyamines, the effects are reversed. This observation can be explained based on the dependence of the interaction between polyamines and DNA on the ionic strength *(33)*: When the ionic strength is reduced, polyamines bind to DNA more strongly, leading to a greater compaction of the chromosome, while an increase in ionic strength leads to the opposite effect. A similar effect as has been observed for polyelectrolyte brushes *(34)*.

#### Fig. S4

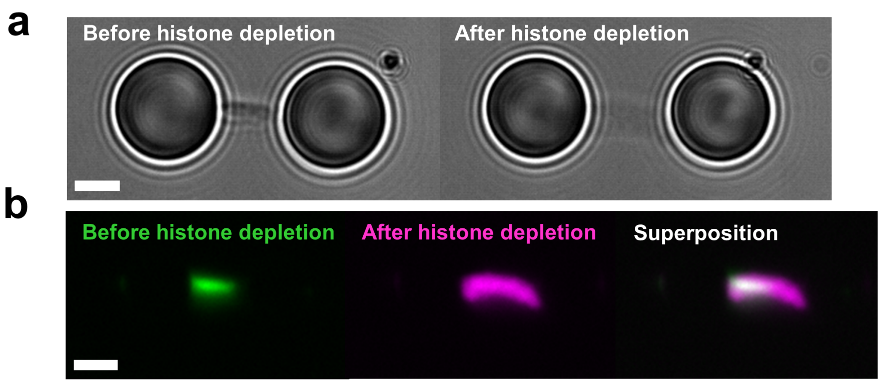


**Figure S4:** (a) Bright field images of a chromosome before and after histone depletion by treatment with 1 M mono valent salt for 15 min. (b) Fluorescence images before and after histone depletion of a chromosome stained with the intercalator SYBR Gold.

### Fig. S5

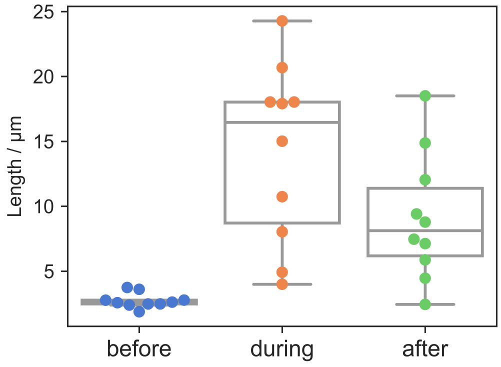


**Figure S5:** Box plot of the Chromosome length before, during and after histone depletion by treatment with 1 M monovalent salt (in the presence of polyamines).

### Fig. S6

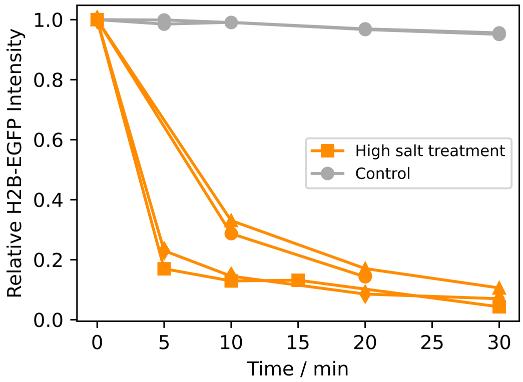


**Figure S6:** Following the depletion of histones by monitoring the fluorescence intensity of EGFP-labelled H2B.

### Fig. S7

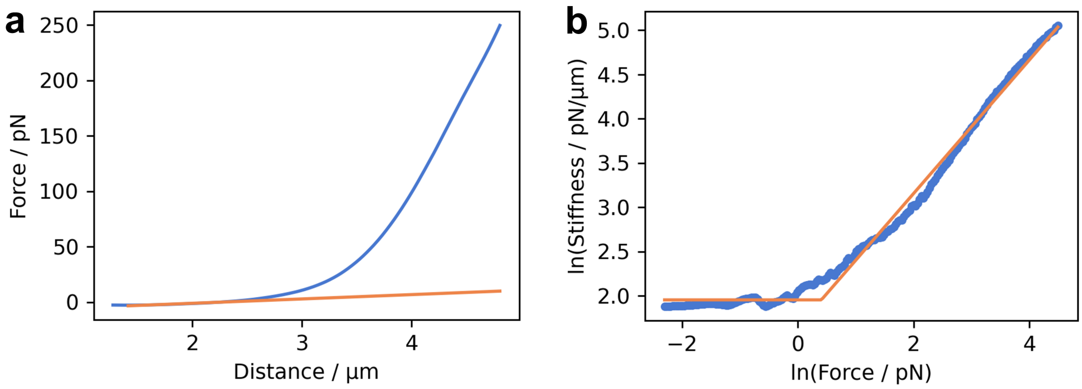


**Figure S7:** Illustration of the parametrization of stretch curves. (a) The linear stiffness is determined as the slope of a linear fit to the initial part of the stretch curve. (b) The critical force is determined by fitting a stepwise function to the stiffness as a function of force. Data is resampled on a logarithmic force scale.

#### Fig. S8

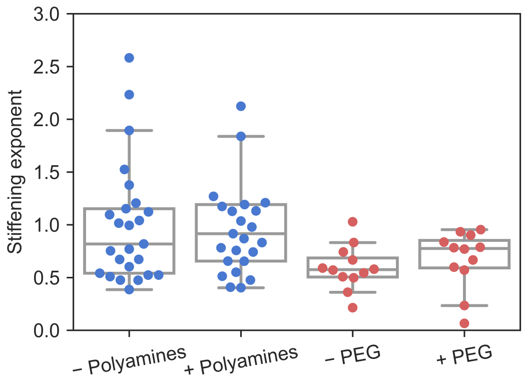


**Figure S8:** Box plot of the stiffening exponent (slope of high force part in Fig. S7B) in the absence and presence of polyamines or PEG confirming that the characteristic non-linear stiffening of chromosomes is not impacted by ion-mediated condensation.

#### Fig. S9

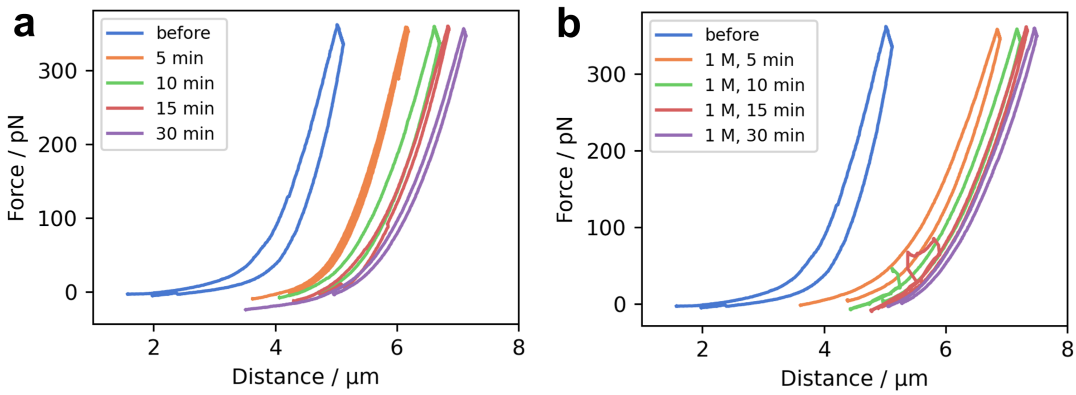


**Figure S9:** Interplay of histone-depletion, ionic strength and polyamines (A) Box plot of the linear stiffness of chromosomes before and after histone depletion and in 1 M salt buffer.(B) Stretch-curves of a chromosome before histone depletion and after varying times of treatment with 1 M salt, recorded in 1 M salt buffer. When chromosomal mechanics were probed in 1 M salt buffer, the lengthening was even more pronounced and independent of the duration of the treatment, since at these high ionic strengths all relevant electrostatic interactions are completely screened. (C) Recorded in regular buffer without polyamines. (D) Recorded in 1 M salt buffer. Interestingly, the behavior of histone depleted chromosomes was again dependent on polyamines. In the presence of polyamines, chromosomes would partially recover when returning from high salt to normal salt conditions, and the lengthening clearly depended on the duration of the high salt treatment. In the absence of polyamines however, the lengthening almost remained as drastic as under high salt conditions. This might indicate that polyamine cations could be able to partially compensate the loss of the positively charged histone complexes, essentially substituting nucleosomes with polyamines.

### Fig. S10

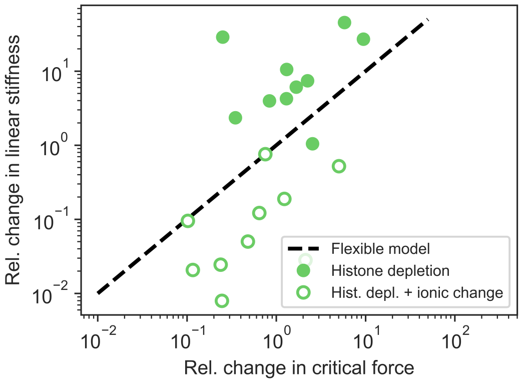


**Figure S10:** Relative change of the linear stiffness as a function of the relative change in critical force for histone depleted chromosomes. The data does not follow a simple scaling relation since the contour length changes drastically during histone depletion.
